# Sex-specific fear acquisition following early life stress is linked to amygdala glutamate metabolism

**DOI:** 10.1101/2024.02.15.580479

**Authors:** Joeri Bordes, Thomas Bajaj, Lucas Miranda, Lotte van Doeselaar, Lea Maria Brix, Sowmya Narayan, Huanqing Yang, Shiladitya Mitra, Veronika Kovarova, Margherita Springer, Karin Kleigrewe, Bertram Müller-Myhsok, Nils C. Gassen, Mathias V. Schmidt

## Abstract

Early life stress (ELS) adversely affects physiological and behavioral outcomes, increasing the vulnerability to stress-related disorders, such as post-traumatic stress disorder (PTSD). PTSD prevalence is significantly higher in women and is partially mediated by genetic risk variants. Understanding how sex influences the interaction of PTSD risk genes, such as *FKBP5*, with trauma-related behaviors is crucial for uncovering PTSD’s neurobiological pathways. The development of in-depth behavioral analysis tools using unsupervised behavioral classification is thereby a crucial tool to increase the understanding of the behavioral outcomes related to stress-induced fear memory formation. The current study investigates the sex-specific effects of ELS exposure by using the limited bedding and nesting (LBN) paradigm. The LBN exposure disrupted different facets of the hypothalamic-pituitary-adrenal (HPA) axis in a sex-specific manner directly after stress and at adult age. Moreover, freezing was altered by LBN exposure in both the acquisition and the retrieval of fear in a sex-dependent manner. Unsupervised behavioral analysis revealed a higher active fear response after LBN exposure during fear acquisition in females, but not in males. The regulation of the HPA axis is closely intertwined with cellular metabolism and core regulatory cascades. To investigate the impact of LBN exposure on tissue-specific metabolism, a metabolomic pathway analysis in the basolateral amygdala revealed a specific sex- and stress-dependent effect on purine, pyrimidine, and glutamate metabolism. The present study highlights the intricate interplay between metabolic pathways and the neurobiological substrates implicated in fear memory formation and stress regulation. Overall, these findings highlight the importance of considering sex-specific metabolic alterations in understanding the neurobiological mechanisms underlying stress-related disorders and offer potential avenues for targeted interventions.

## Introduction

Early life stress (ELS) exposure, such as child abuse or neglect, has severe, long-lasting negative behavioral and physiological consequences in adulthood, including, among others, an altered neuroendocrine function ^1–3^ and morphological changes in the brain ^4,5^. This ultimately leads to an increased risk and persistence of stress-related disorders, such as post-traumatic stress disorder (PTSD) ^6–8^. Human genetic studies have identified that stress-related disorders are partially mediated by different genomic variations ^9–15^, in particular the glucocorticoid receptor (GR) co-chaperone FK506 binding protein 51 kDa (FKBP51), encoded by the FKBP5 gene ^16–22^. An additional important factor influencing the susceptibility and severity of PTSD is sex. The prevalence of PTSD is around twice as high in women (11.0%) compared to men (5.4%) ^23^. However, the increased risk for PTSD development in women could not be entirely explained by differences in the event type or severity of the traumatic event, which might indicate a potential underlying biological mechanism ^24,25^. Therefore, investigating the underlying mechanisms related to ELS exposure in a sex-specific way is crucial to advance the understanding of the neurobiological mechanism of PTSD.

An increasing number of clinical studies point to corticolimbic structures as strongly affected in size and functioning by ELS exposure, in particular the basolateral amygdala (BLA) and hippocampus (HIP) ^26–30^. Moreover, the BLA and HIP continue their development in function and morphology during the early postnatal period ^31–33^, rendering this developmental period especially vulnerable to environmental insults. Furthermore, both the BLA and HIP exhibit high expression levels of GRs and mineralocorticoid receptors (MRs) ^34–37^, making them particularly vulnerable to the effects of ELS exposure. Animal models have further elucidated a significant role of FKBP5 in these regions, demonstrating a high expression under baseline conditions, and a strong upregulation following stress ^38^.

Alterations of anxiety and fear behavior are a central hallmark of a PTSD-like phenotype in animal models. A study using a mouse model, overexpressing the human FKBP5 gene in the forebrain showed that elevated FKBP5 expression in combination with ELS exposure increases anxiety-related behavior, which was more pronounced in females ^39^. Therefore, the genetic regulation of FKBP5 is linked to affect males and females differently after the exposure to ELS ^40–42^, but the impact of ELS exposure on FKBP5 regulation remains to be investigated. There is an increasing body of evidence that ELS exposure affects rodents in a sex-specific manner ^43–47^. However, the effects of fear memory formation on ELS have not been explored in detail. Exposure to fear conditioning in rodents has been shown to strongly activate the BLA and HIP in a time-dependent manner ^48^. Therefore, the important role of the BLA and HIP in ELS exposure indicates that the formation of fear acquisition and memory could be affected by ELS exposure. Previous research has shown that ELS reduces fear expression during contextual as well as auditory fear memory retrieval in males, which is linked to a reduction of synaptic plasticity markers in the dorsal HIP ^49^, but the sex-dependent effects and exact behavioral mechanisms remain to be uncovered.

The exploration of metabolic mechanisms and pathways in the brain are an important assay contributing to the understanding of the underlying neurobiological mechanisms related to the sex-dependent effects of ELS exposure. Previous research has shown the importance of the metabolic purine and pyrimidine pathways in the HIP when looking at successful treatment response of chronic intervention with the selective serotonin reuptake inhibitor paroxetine ^50^. However, the exploration of metabolism and the underlying regulatory pathways in specific stress-related and affected brain regions in relation to ELS has remained uninvestigated.

In the current study, we address these open questions by investigating the sex-specific effects of ELS exposure using the established limited bedding and nesting (LBN) paradigm on FKBP5 expression, brain tissue metabolomics, and fear acquisition and retrieval. Utilizing an unsupervised deep phenotyping strategy, we show that specific aspects of fear behavior and memory are altered by ELS in a sex-dependent manner. These behavioral alterations align with FKBP5 expression changes in BLA and HIP. Moreover, metabolic pathway analysis in the BLA revealed a sex- and stress-dependent effect on purine, pyrimidine, and glutamate metabolism. The present study highlights the intricate interplay between stress-responsive genes, metabolic pathways, and the neurobiological substrates implicated in fear memory formation and stress regulation.

## Methods

### Animals

Adult male and female C57/Bl6N mice (age between 2-3 months of age) were obtained from the in-house facility of the Max Planck Institute of Psychiatry and used for breeding (F_0_). Animals from the F_1_ generation were used as experimental animals and were weaned at P25 in groups of maximum four animals with their littermates. Animals were housed in individually-ventilated cages (IVC; 30cm×16cm×16cm connected by a central airflow system: Tecniplast, IVC Green Line—GM500). All animals were kept under standard housing conditions; 12h/12h light-dark cycle (lights on at 7 a.m.), temperature 23±1°C, humidity 55%. Food (Altromin 1324, Altromin GmbH, Germany) and tap water were available ad libitum. All experimental procedures were approved by the committee for the Care and Use of Laboratory Animals of the government of Upper Bavaria, Germany. All experiments were in accordance with the European Communities Council Directive 2010/63/EU.

### ELS paradigm: limited bedding and nesting

ELS was performed using the LBN paradigm to induce chronic stress towards the mother and pups during P02 to P09, as previously described by Rice et al. ^51^. At P02, all litters were transferred to new IVCs and randomly assigned to the non-stressed (NS) or stressed (LBN) condition. If necessary, the litters were culled to a maximum of 10 animals per litter. The LBN litters were placed on a stainless-steel mesh (McNichols) and were provided with limited nesting material (1/2 square of Nestlets, Indulab). The NS animals were placed in an IVC with a standard amount of bedding material and were provided with a sufficient amount of nesting material (2 squares of Nestlets). All litters were left undisturbed until P09, after which they returned to standard housing conditions. The pups were weaned in same-sex groups with a maximum of four animals per cage.

### Adult behavioral testing

At 3 months of age, a cohort of both males and females, were tested on a fear conditioning protocol containing fear acquisition (day 1) and the subsequent recall of contextual fear memory (day 2) and auditory fear memory (day 3). The behavioral tests were performed between 8 a.m. and 11 a.m. in the same room as the housing facility.

### Fear conditioning

The fear conditioning protocol was performed as previously described ^52^. Data were recorded and analyzed using the ANY-maze 7.2 software (Stoelting, Ireland), in which the percentage of the time freezing was calculated. Furthermore, the fear acquisition data were subsequently analyzed using DeepLabCut version 2.2b7 ^53^ and DeepOpenField (DeepOF) version 0.2 ^54,55^ for the unsupervised analysis pipeline.

#### Fear acquisition

The fear acquisition consisted of placing the mice into a cube-shaped fear conditioning chamber (Bioseb, France) with a metal grid floor to provide electric shocks. At the start of the test, the chamber light was switched on, and after an initial habituation time of 3 min, the mice were exposed to five conditioned-unconditioned stimulus pairings (auditory conditioning stimulus: 30 sec, 9kHz, 80dB tone & unconditioned stimulus 0.5 sec, 0.6mA foot shock) with an inter-trial interval (ITI) of 5 mins. 1 min after the last foot shock the animals were returned to their home cage. Before and after each trial, the conditioning chamber was thoroughly cleaned with 70% EtOH. The calculation of the mean freezing statistics was performed using the average of tones 2-5, leaving out the first tone, as no shock history was present at that moment. The mean ITI freezing was calculated using all four ITIs.

#### Contextual fear memory

Contextual fear memory was performed 24 hours after initial fear acquisition. The same setup was used as by fear acquisition, except that no conditioned-unconditioned stimulus protocol was executed. The test endured for a total of 5min in which only the chamber light was switched on, and again before and after each trial, the conditioning chamber was thoroughly cleaned with 70% EtOH.

#### Auditory fear memory

The consolidation of auditory fear memory was performed two days after the fear acquisition. The set-up was replaced by a novel and neutral context, which differed in material (plexiglass), shape (circular), and surface texture, as no grid was present at the bottom of the set-up. In addition, the cleaning solution was changed in odor, using 1% acetic acid. This allowed for measuring the fear response specifically towards the tones, without the interference of the context. The chamber light was switched on at the start of the test, after which mice were left undisturbed for an initial 1 min habituation phase. Then, the mice were exposed to the same tones as heard in the fear acquisition (30 sec, 9kHz, 80dB) 15 times with a 1.5 min ITI. 1 min after the last tone the animals were returned to their home cage. The mean tone and ITI freezing were calculated using all tones and ITIs.

### Unsupervised analysis of the fear conditioning data

An additional unsupervised analysis was performed for the fear acquisition data in both males and females in order to obtain a more in-depth analysis of the behavioral differences between conditions and sexes during the ITIs and tones 2-5. First, pose estimation was performed on the raw data videos using DeepLabCut version 2.2b7 (single animal mode). DeepLabCut pose estimation analysis was performed using 11 body parts, including the nose, left and right ears, three points along the spine (including the center of the animal), all four extremities, and the tail base.

Subsequently, DeepLabCut annotated datasets were processed and analyzed using DeepOF v0.2, as described previously ^54,55^. In brief, DeepOF preprocesses the DeepLabCut annotated data by performing rotational alignment and centering of the coordinates. The unsupervised analysis of the fear acquisition data was performed on the entire video length and utilized the same model (DeepOF - Variational Deep Embeddings (VaDE)) for the male and female data in order to make cluster interpretation between sexes possible. The interpretation of the clusters was explored by visual inspection of representative video snippets for each specific cluster. Representatives were selected as instances with a cluster assignment confidence greater or equal than 0.9. In addition, Shapley additive explanations (SHAP) were utilized to rank feature importance per cluster in order to further understand the expressed behavior within clusters.

### Physiological measurements

One week following the behavioral tests, adult animals were weighed and subsequently sacrificed by decapitation, after which trunk blood was collected in EDTA-coated microcentrifuge tubes (Kabe Labortechnik, Germany) and directly transferred to ice. Samples were centrifuged at 4°C for 15min at 8.000 rpm, after which plasma was removed and kept transferred for storage at - 80°C. A separate cohort of mice was used to obtain corticosterone (CORT) measurements directly after the stress at P09. On the morning of P09, litters were kept in their cage, while first the mother was sacrificed, and subsequently the pups were sacrificed, keeping them in their nest as long as possible to minimize the influence of acute stress exposure. Trunk blood was collected and processed as described for adult blood samples. Plasma CORT levels were measured in duplicates using radioimmunoassay following the manufacturer’s protocol (MP Biomedicals, Eschwege, Germany). Adrenals were dissected and kept at 4°C in saline (0.9% NaCl) until further processing, which included the removal of all surrounding fat tissue and weighing. The relative adrenal weight was calculated by dividing the total body weight before sacrifice by the total adrenal weight, including the adrenals from both sides.

### In-situ hybridization of FKBP5

The FKBP5 mRNA profile was determined using radio-active *in-situ* hybridization labeling as described previously ^38^. In brief, the animals were either sacrificed directly after the stress exposure at P09 or in adulthood at 2 months of age. After decapitation, the brains were removed and snap-frozen using 2-methyl butane (kept on dry ice) and stored at -80°C until further use. Brains were sliced using a cryostat in 20 µm sagittal sections, which resulted in a series of the BLA and dorsal HIP slides that were thaw-mounted on Super Frost Plus Slides and stored at -20°C. The *in-situ* hybridization sections were removed from -20°C, left to dry at room temperature, fixated with 4% paraformaldehyde, and subsequently dehydrated using a series of increasing concentrations of ethanol. Then, the hybridization buffer was equally spread out over the different slides containing the radioactive ^35^S-UTP-labeled FKBP5 riboprobe and incubated overnight at 55°C. On the next day, the sections were rinsed, incubated with RNAse A, desalted, and dehydrated, after which the radioactive slides were exposed to Kodak Biomax MR films (Eastman Kodak Co., Rochester, NY) and developed after an exposure time of 12 days. Films were digitized, and the regions of interest were identified using the mouse brain atlas (https://developingmouse.brain-map.org/static/atlas). The expression was determined by optical densitometry with the ImageJ software (NIH, Bethesda, MD, USA). The expression was averaged per brain region per animal and subtracted by the background signal of a nearby structure that did not express the *fkbp5* gene. A distinction was made between important subregions of the dorsal HIP, in which separate measurements were obtained for the CA1, CA2-3, and the dentate gyrus (DG).

### Metabolomics

#### Extraction of polar metabolites from mouse BLA tissue samples

To extract polar metabolites from mouse (BLA tissue samples, a sample extraction buffer consisting of methyl tert-butyl ether (MTBE, Sigma-Aldrich, 650560), methanol (Carl Roth, P717.1), and water (Biosolve, 232141), all of LC-MS grade, in a volumetric ratio of 50:30:20 [v:v:v] was used. The sample extraction buffer contained the following internal standards: U-13C15N-labeled amino acids at a final concentration of 0.25 µM (prepared as 2.5 mM in 0.1 N HCl, Cambridge Isotope Laboratories, MSK-A2-1.2), citric acid d4 at 0.02 µg/mL (dissolved at 100 µg/mL in H2O, Sigma-Aldrich, 485438-1G), ATP 13C10 at 0.1 µg/mL (dissolved at 1 mg/mL in 5 mM Tris-HCl, Sigma-Aldrich, 710695), AMP 13C10, 15N5 at 0.1 µg/mL (dissolved at 1 mg/mL in 5 mM Tris-HCl, Sigma-Aldrich, 650676), ADP 15N5 at 0.1 µg/mL (dissolved at 1 mg/mL in 5 mM Tris-HCl, Sigma-Aldrich, 741167), and EquiSPLASH™ LIPIDOMIX at 0.02 µg/mL (prepared as 100 µg/mL, Avanti Polar Lipids, 30731).

The sample extraction buffer was freshly prepared and cooled to -20°C. Subsequently, 1 mL of the chilled sample extraction buffer was added to the homogenized tissue, which had been preprocessed with 5 mm metal balls in a TissueLyser (Qiagen, TissueLyser LT) for 1 minute at 25 Hz. The mixture was then incubated at 4°C for 30 minutes with agitation at 1500 rpm using a thermomixer (VWR, Thermal Shake lite).

Following incubation, metal balls were removed, and the samples underwent centrifugation (Thermo Scientific, Fresco 21) at 4°C for 10 minutes at 21,000 x g, resulting in the transfer of the cleared supernatant to a 2 mL tube. To this supernatant, 200 µL of MTBE and 150 µL of water were added, followed by incubation at 15°C for 10 minutes with agitation at 1500 rpm in a thermomixer. A subsequent centrifugation at 15°C for 10 minutes at 16,000 x g facilitated phase separation. Approximately 650 µL of the upper lipid-containing phase was transferred to a separate 1.5 mL tube (not included in this study). The remaining polar metabolite extract, approximately 600 µL in volume after removal of the residual lipid phase, was dried using a SpeedVac concentrator (Eppendorf, 5301 Vacufuge).

The resulting dried metabolite pellets were subsequently stored at -80°C until further analysis.

#### HILIC-MS for profiling of metabolites in mouse BLA tissue

The samples were dissolved using 100 µL of 70% MeOH (v/v). After 10 min of shaking at 1100 rpm at 10°C with a ThermoMixer® C (Eppendorf AG, Hamburg, Germany), the samples were centrifuged at 13 000 rpm for 10 min at 10°C with Centrifuge 5424 R (Eppendorf AG, Hamburg, Germany). The clear supernatant was transferred to a 1.5 mL glass vial with insert. A QC-sample was pooled by combining 5 µL of each sample.

The untargeted analysis was performed using a Nexera UHPLC system (Shimadzu, Duisburg, Germany) coupled to a Q-TOF mass spectrometer (TripleTOF 6600 AB Sciex, Darmstadt, Germany). Separation of the samples was performed using a UPLC Premier Amide 2.1 × 100 mm, 1.7 µm analytic column (Waters, Eschborn, Germany) with a 400 µL/min flow rate. The mobile phase was 5 mM ammonium acetate in water (eluent A) and 5 mM ammonium acetate in acetonitrile/water (95/5, v/v) (eluent B). The gradient profile was 100% B from 0 to 1.5 min, 60% B at 8 min and 20% B at 10 min to 11.5 min and 100% B at 12 to 15 min. A volume of 5 µL per sample was injected. The autosampler was cooled to 10 °C and the column oven heated to 40 °C. The samples were measured in a randomized order and in the Information Dependent Acquisition (IDA) mode. MS settings in the positive mode were as follows: Gas 1 55, Gas 2 65, Curtain gas 35, Temperature 500 °C, Ion Spray Voltage 5500, declustering potential 80. The mass range of the TOF MS and MS/MS scans were 50–2000 m/z and the collision energy was ramped from 15– 55 V. MS settings in the negative mode were as follows: Gas 1 55, Gas 2 65, Cur 35, Temperature 500 °C, Ion Spray Voltage –4500, declustering potential –80. The mass range of the TOF MS and MS/MS scans were 50–2000 m/z and the collision energy was ramped from –15–55 V.

The “msconvert” from ProteoWizard ^56^ was used to convert raw files to mzXML (de-noised by centroid peaks). The bioconductor/R package xcms ^57^ was used for data processing and feature identification. More specifically, the matchedFilter algorithm was used to identify peaks (full width at half maximum set to 7.5 seconds). Then the peaks were grouped into features using the “peak density” method ^57^. The area under the peak was integrated to represent the abundance of features. The retention time was adjusted based on the peak groups present in most samples. To annotate features with names of metabolites, the exact mass and MS2 fragmentation pattern of the measured features were compared to the records in HMBD ^58^ and the public MS/MS spectra in MSDIAL ^59^, referred to as MS1 and MS2 annotation and their standards, respectively. Missing values were imputed with half of the limit of detection (LOD) methods, i.e., for every feature, the missing values were replaced with half of the minimal measured value of that feature in all measurements. To confirm a MS2 spectra is well annotated, we manually reviewed our MS2 fragmentation pattern and compared it with records in the public database or previously measured reference standards as well as to SIRIUS ^60^ to evaluate the correctness of the annotation.

Annotated features were subjected to a metabolite set enrichment analyses, MSEA, using KEGG pathway-based library ^61^ with the only use of metabolite sets containing at least 2 entries in MetaboAnalyst 5.0 ^62–65^. The metabolic analysis investigates the expression of a set of metabolites using two different settings, including a HILIC negative mode and a HILIC positive mode. These different HILIC settings can reveal partially different metabolites, and are merged to obtain complete list of metabolites.

### Statistics

Statistical analyses and graphs were made using RStudio (with R 4.1.1), except for the unsupervised DeepOF analysis, which was performed using Python (v 3.9.13). Different batches of animals were used for the adult fear conditioning behavior (female: LBN n=10, NS n=10 & male: LBN n=11, NS n=11) and the in-situ hybridization experiments (female P09: LBN n=4, NS n=6 & male P09: LBN n=5, NS n=3, female adult (2 months): LBN n=5, NS n=4 & male adult (2 months): LBN n=3, NS n=4). All animals were used for statistical analysis unless stated otherwise. During the contextual fear memory, 2 NS male animals were excluded from the analysis due to technical difficulties. Data were tested for the corresponding statistical assumptions, which included the Shapiro-Wilk test for normality and Levene’s test for heteroscedasticity. If assumptions were violated the data were analyzed using non-parametric variants of the test. The group comparisons were analyzed using the independent samples t-test (T) as a parametric option, Welch’s test (We), if data was normalized but heteroscedastic, or the Wilcoxon test (Wx) as a non-parametric option. The time-binned data (fear conditioning) was analyzed using the two-way repeated measures ANOVA with the phase (e.g. tones) as a within-subject factor and the condition (non-stressed vs. stressed) as a between-subject factor. Data that showed a significant main effect were further analyzed with the post-hoc Bonferroni test (parametric) or the Kruskal-Wallis test (non-parametric). P-values were adjusted for multiple testing using the Bonferroni method. The timeline and bar graphs are presented as mean ± standard error of the mean (SEM). Data were considered significant at p<0.05 (*), and further significance was represented as p<0.01 (**), p<0.001 (***), and p<0.0001 (****).

## Results

### The physiological hallmarks of limited bedding and nesting (LBN) exposure

Chronic stress during early life was induced to assess the sex-dependent stress effects directly after early stress exposure and in adult age (figure 1A). A common consequence of LBN exposure is the reduction in body weight at P09 ^51^. The current study observed a significant reduced body weight in males and females, demonstrating a successful induction of chronic stress exposure during early life (figure 1B). Interestingly, the stress-induced reduction in body weight was sustained in female adult age, but not in males (figure 1C). A marker for chronic stress exposure and dysregulation of the HPA axis is the relative weight of the adrenals, which in adult age was not altered in females, but was significantly elevated in LBN males (figure 1D). In addition, as a proxy for stress exposure, the CORT levels were obtained directly after the LBN exposure at P09, the female LBN animals showed a significant elevation of CORT levels, whereas in males no elevated CORT levels were observed (figure 1E). In adult age, no difference in basal CORT levels was observed for stress exposure in both sexes (figure 1F).

**Figure 1.**
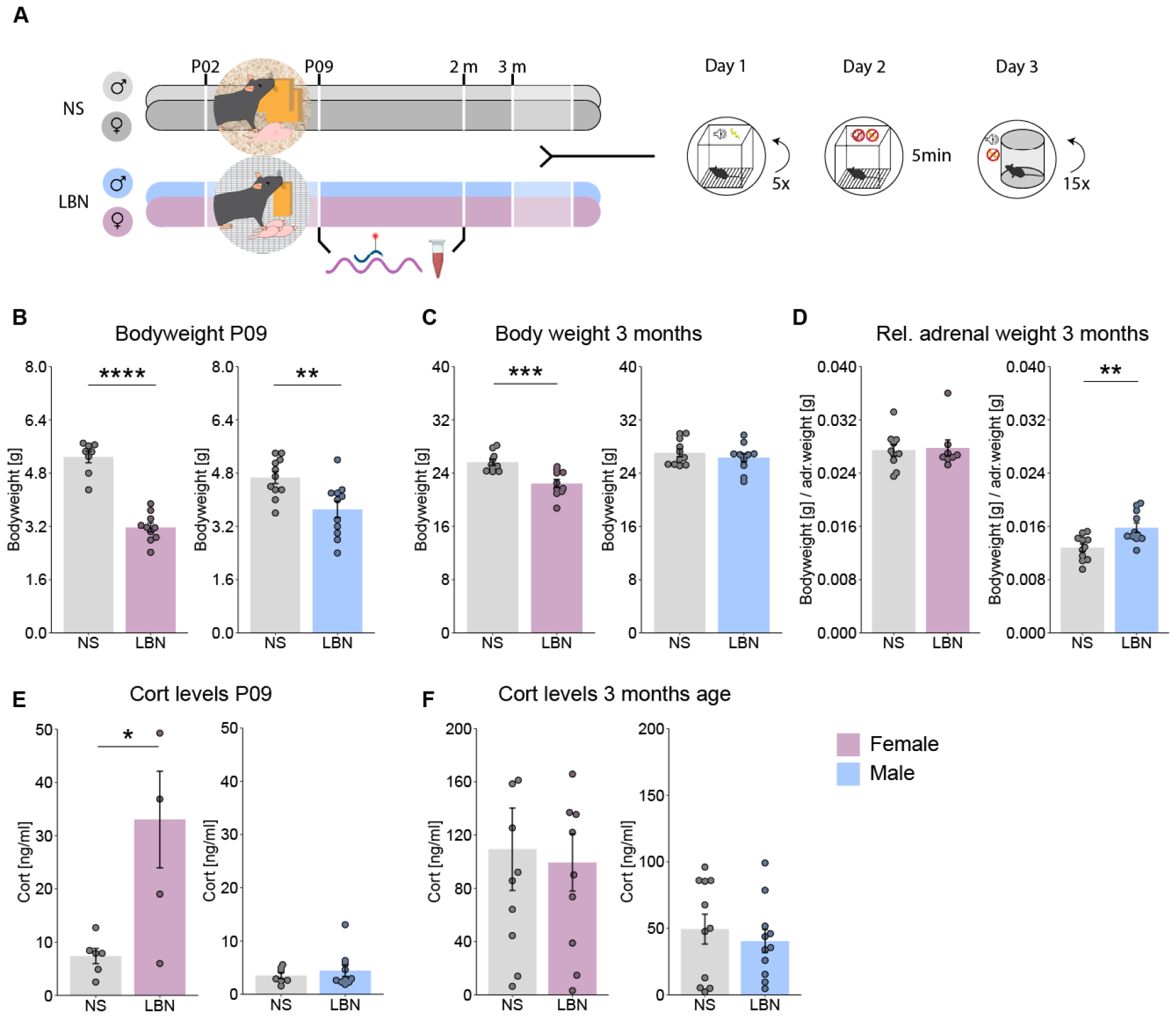
Physiological stress hallmarks of limited bedding and nesting (LBN). **A)** Experimental timeline for non-stressed (NS) controls and limited bedding and nesting (LBN) stressed mice on fear conditioning. **B)** Significant decrease in body weight was observed after LBN exposure at P09 for females (Wx=80, p<0.0001), and males (T(20)=3.13, p=0.005). **C)** During adult age (3 months) the body weight was significantly reduced in females (T(18)=4.16, p=0.0006), but not in males (T(20)=0.88, p=0.39). **D**) The relative adrenal weight was not altered in females (Wx=45, p=0.70), but was significantly increased in LBN males (T(20)=-3.3, p=0.003). **E)** At P09, CORT levels were significantly elevated in LBN females (Wx=4, p=0.05), whereas this was not the case for males (Wx=31, p=0.74). **F)** At adult age, the CORT levels were not altered by stress for both females (T(18)=0.26, p=0.79) and males (T(20)=0.64, p=0.53). The bar graphs are presented as mean ± standard error of the mean and all individual samples as points. Panels B-D, F represent female NS (n=10), female LBN (n=10), male NS (n=11), male LBN (n=11). Panel E represent female NS (n=6), female LBN (n=5), male NS (n=7), male LBN (n=10).

### LBN increased FKBP5 expression in the dorsal hippocampus exclusively in males

FKBP5 mRNA expression in the BLA (supplemental figure 1A) and dorsal HIP, separating the subregions; CA1, CA2-3, and DG (supplemental figure 1B) was assessed directly after LBN exposure at P09 and in adult age. Female and male data at P09 did not show a stress-induced difference of FKBP5 expression in the BLA, CA1, CA2-3, and DG (supplemental figure 1C, D), however, an indication of elevated FKBP5 expression in LBN animals could be observed in males, but this was not significant (supplemental figure 1D). Moreover, also during adult age, the female data did not indicate any FKBP5 expression differences between stress conditions (supplemental figure 2E). However, adult males showed a significant stress-induced increase of FKBP5 expression in the CA1 region, but not in the BLA, CA2-3, and DG regions (supplemental figure 1F). Moreover, both NS and LBN mice showed an age-dependent expression pattern of FKBP5 regardless of sex in the dorsal HIP. At P09 FKBP5 expression was the highest in the CA2-3 and showed similar levels of expression in the CA1 and DG, whereas at p56, the FKBP5 expression was the highest in the DG, followed by the CA2-3, and the CA1 (supplemental figure 2C-F).

**Figure 2.**
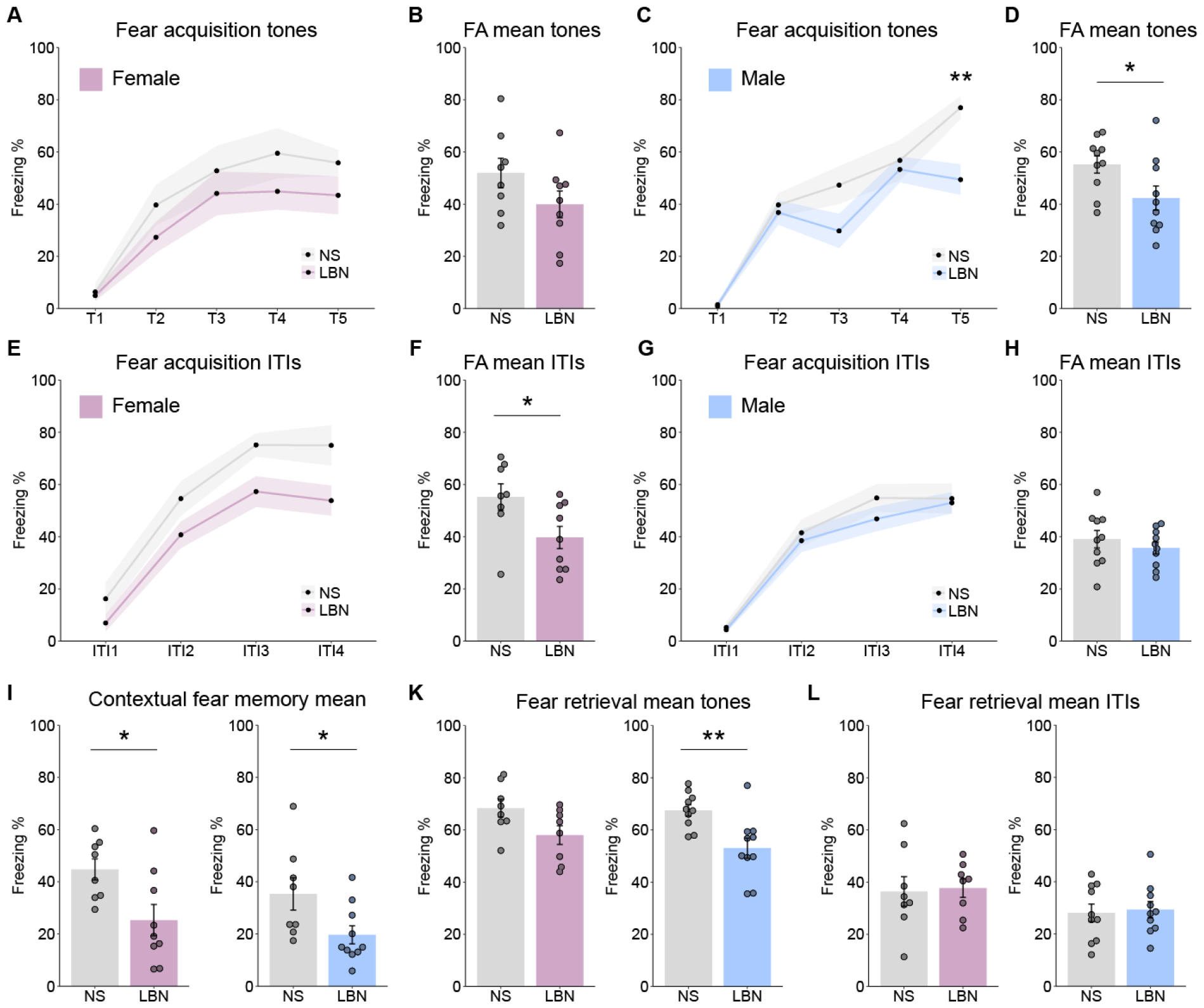
Fear conditioning data on freezing behavior. **A)** Freezing behavior on individual tones during fear acquisition (FA) was not altered between non-stressed (NS) and stressed (LBN) females, two-way ANOVA (p>0.45). **B)** The average freezing during the FA tones 2-5 was not altered between NS and LBN females (T(15)=1.58, p=0.14). **C)** Freezing behavior in the FA for individual tones 1-4 was not altered between NS and LBN males (p>0.22), but for tone 5 it was significantly lowered in LBN males (F(1,18)=14.3, p=0.005), with a significant main effect for the two-way ANOVA on stress (F(1,90)=9.39, p=0.003), tones (F(4,90)=41.30, p<0.0001), and stress*tones (F(4,90)=2.6, p=0.041). **D)** The average freezing during FA tones 2-5 was significantly lowered in LBN males compared to NS (T(18)=2.25, p=0.037). **E)** No significant main effect could be observed between NS and LBN females for the individual ITIs in the FA. **F)** The average freezing during FA ITIs 1-4 was significantly lowered in LBN females compared to NS (T(15)=2.37, p=0.03). **G)** No significant main effect could be observed between NS and LBN males for the individual ITIs in the FA. **H)** The average freezing during FA ITIs 1-4 was not altered between NS and LBN males (T(18)=0.84, p=0.41). **I)** The mean freezing in the contextual fear memory task was significantly lowered in LBN females compared to NS (T(15)=2.61, p=0.020), and in males (T(16)=2.32, p=0.034). **K)** The mean freezing in the fear retrieval task showed no significant difference in females (T(14)=2.09, p=0.055). However, in males, a significant reduction in freezing was observed in LBN compared to NS (T(18)=3.29, p=0.004). **L)** No significant differences were observed in the mean freezing during the ITIs between LBN and NS animals for females and males. The timelines and bar graphs are presented as mean ± standard error of the mean and all individual samples as points. Panel A-L represent female NS (n=10, female LBN (n=10), male NS (n=10), male LBN (n=11).

### Freezing behavior is affected by LBN exposure in a sex-specific manner

During the acquisition of fear conditioning, the typical increase of freezing behavior over the different tone representations was observed in both females and males, regardless of the stress condition (figure 2A-D). However, an overall decrease during the fear acquisition in freezing behavior was observed at the tone representations in males, but not females (figure 2B, D). Moreover, the exploration during the ITIs of the fear acquisition phase showed that the freezing at the individual ITIs was not significantly altered in females based on the stress condition (figure 2E), but there was a significant reduction in the freezing response of LBN females in the overall mean ITIs (figure 2F), which was not observed in males (figure 2G-H). When looking at the recall of fear, the mean freezing during the contextual fear memory was significantly lowered in LBN females and males (figure 2I, supplemental figure 2A-B). The auditory fear retrieval did not show a different freezing response on mean tones in females (figure 2K, supplemental figure 2C), but did show a lowered freezing response in LBN males compared to NS (figure 2K, supplemental figure 2D). In addition, no differences were observed in the fear retrieval for both females and males (figure 2L, supplemental figure 2E-F).

The DeepOF unsupervised clustering analysis of the fear acquisition data yielded 9 distinct clusters (figure 3A, B). No cluster population differences were observed by the stress background for both female and male ITI 1-4 data (supplemental figure 3 A, B). However, clusters were significantly altered by the stress background in females during tones 2 to 5 (figure 3A), which were not observed in males (figure 3B). In particular, “cluster 0” was significantly increased in LBN females, and “cluster 6” significantly decreased in LBN females compared to NS (figure 3A). A multi-class supervised learning model was trained to map from motion summary statistics to the obtained cluster labels, and performance was measured in terms of the balanced accuracy per cluster (figure 3C). The confusion matrix showed low probabilities for all cluster crossovers, and the classifier performance was substantially greater than random for all clusters, indicating that all clusters were substantially distinguishable by the model (figure 3C). The cluster detection analysis yields a set of feature explainers per cluster that can be used to interpret the clusters using SHAP values in global (figure 3D) and cluster-specific ways (figure 3E, F). Importantly, the interpretation of the clusters was done using the feature importance of the SHAP analysis, together with the visual interpretation of the video fragments per cluster (see supplemental materials for video output per cluster). The global feature importance across all clusters revealed that the distance towards several spine labels (a stretch or a shortening of the back), the overall speed (an increased or reduced speed), huddle (an increased or decreased amount of the behavior in which the animal stops moving around and bends the back), and the surface area of the head (an increased surface area is related to the head being forward, whereas a decreased surface area is related to the head being downward) were particularly important for global cluster inclusion (figure 3D). More specifically, the feature importance analysis for “cluster 0” revealed that an increased speed, a decrease in huddle behavior, and an increased spine stretch were important features for cluster inclusion (figure 3E). The visual inspection of “cluster 0” indicated a behavior related to the exploration of the environment, in which the animal was moving around (see supplemental materials). In contrast, “cluster 6” feature importance analysis revealed that a decrease in spine stretch, an increase in huddle behavior, and a reduction of the surface head area were important features for cluster inclusion (figure 3F). The visual inspection of “cluster 6” indicated a behavior related to freezing, in which the animal was often immobile and close to the outside of the environment.

**Figure 3.**
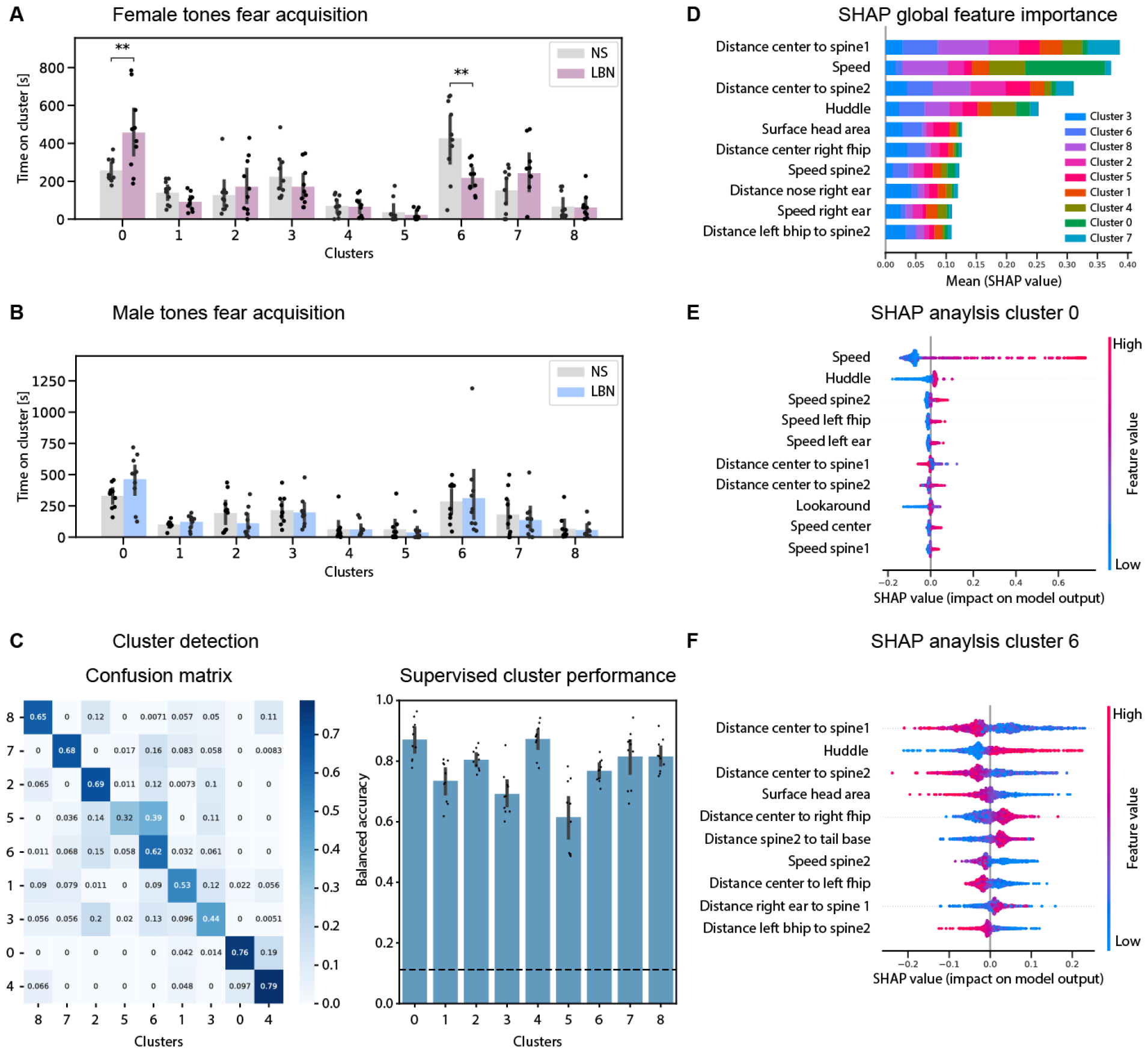
Unsupervised analysis of fear acquisition data on tones 2-5. **A)** Cluster enrichment for the female fear acquisition data using tones 2-5. Bar graphs represent mean ± standard deviation of the time proportion spent on each cluster. Statistics were performed using an independent samples t-test corrected for multiple testing using Benjamini-Hochberg’s method across clusters between NS and LBN exposure. Significant differences were observed in clusters 0: T=-3.55, p=2.28^e-3^, and 6: T=4.24, p=4.97^e-4^, but none of the other clusters (p>0.05). **B)** Cluster enrichment for the male fear acquisition data using tones 2-5. No significant differences were observed between NS and LBN mice using the independent samples t-test corrected for multiple testing using Benjamini-Hochberg’s method across clusters (p>0.05). **C)** On the left, the confusion matrix is obtained from the trained gradient boosting machine classifying between clusters. Aggregated performance over the validation folds of 10-fold cross-validation is shown. On the right, is the validation performance per cluster across a 10-fold cross-validation loop. Balanced accuracy was used to correct for cluster assignment imbalance. The dashed line marks are the expected performance due to chance, considering all outputs. **D)** The global SHAP feature importance between the different clusters. Features in the y-axis are sorted on the global absolute SHAP values across all clusters. The classes in the bar graphs are sorted by highest to lowest clusters importance within every feature. **E-F)** Bee swarm plots for the two differentially expressed clusters within the female fear acquisition data between NS and LBN mice, clusters 0 and 6. The plots show the 10 most important features for each classifier, in terms of the mean absolute value of the SHAP values. Bar graphs represent mean ± standard deviation of the time proportion spent on each cluster. Panel A-F represent female NS (n=10, female LBN (n=10), male NS (n=10), male LBN (n=11).

### Sex-dependent LBN exposure affect the expression of metabolic pathways in the BLA

In order to determine the intertwined relation between sex-specific and brain region-specific stress effects with global metabolic cascades, a comprehensive and highly sensitive metabolomic measurement was conducted. The metabolomic analysis revealed a significant regulation (down or up) for sex, by comparing males and females regardless of stress exposure in HILIC negative mode (62 significant features, p<0.05) and positive mode (105 significant features) (supplemental figure 4A). The regulation through stress did not reveal a pronounced regulation of metabolic cascades, comparing NS and LBN conditions, regardless of sex in HILIC negative mode (0 significant features) and positive mode (2 significant features) (supplemental figure 4A).

**Figure 4.**
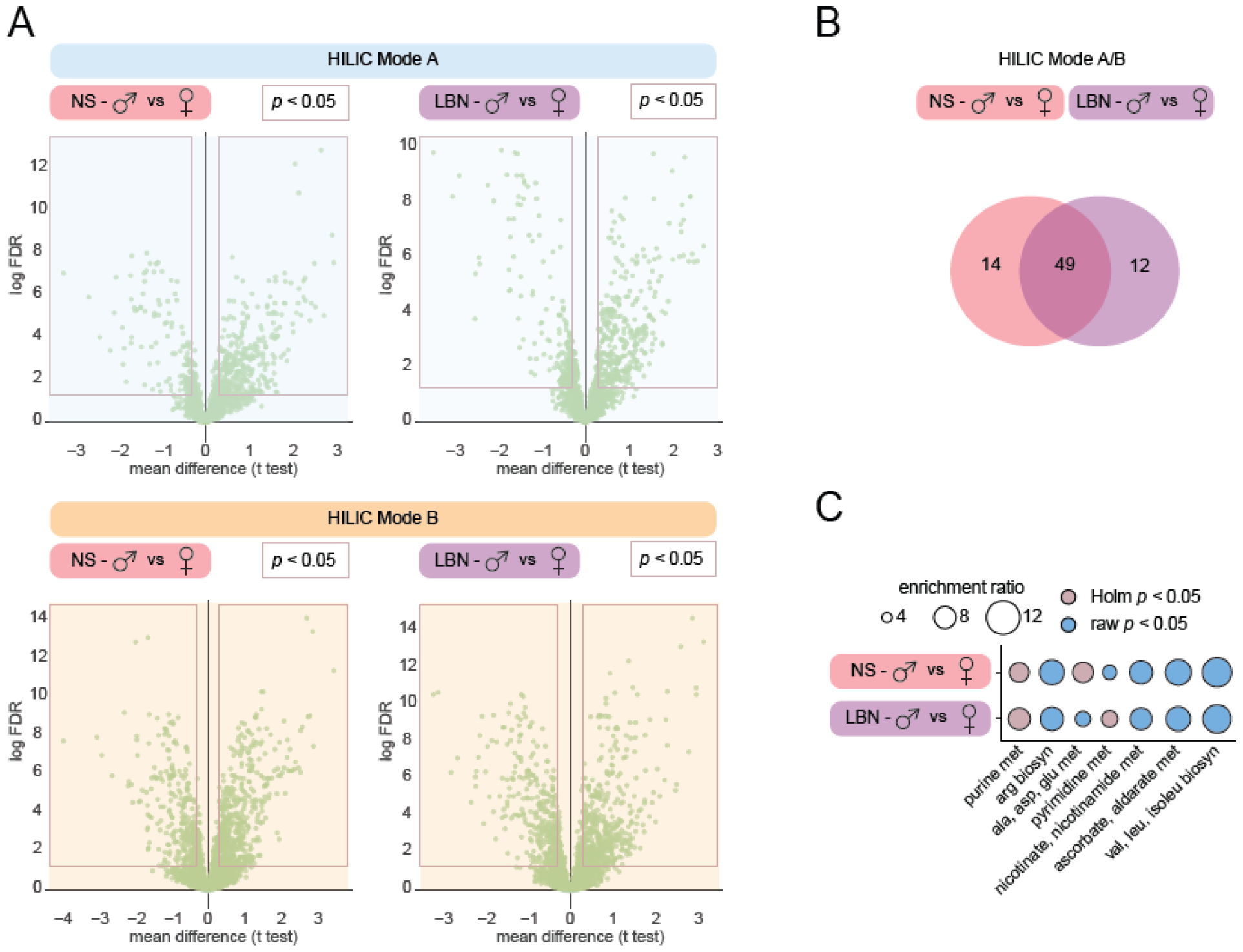
Metabolomics reveals altered glutamate, pyrimidine, and purine pathways related to sex and stress condition. **A)** Volcano plots show a large differential expression profile comparing male vs females in NS, as well as LBN conditions in HILIC mode A and B. **B)** The set of regulated metabolites in the male vs female comparison between NS and LBN conditions showed that 34.67% of the metabolites were specifically regulated in only NS or only LBN conditions (18.67%: 14 metabolites in NS and 16%: 12 metabolites in LBN of the total 75 metabolites) **C)** Metabolite Set Enrichment Analysis (MSEA) revealed different pathways of interest. 1) purine metabolism was significantly regulated in NS (Holm test; p=1.28^e-6^; enrichment ratio (ER)=7.32) and LBN (Holm test; p= 1.43^e-8^; ER=8.09). 2) pyrimidine metabolism was significantly regulated in LBN (Holm test; p=0.021; ER=6.23), but not in NS (Holm test; p=0.13; raw p-value=0.0017; ER=5.48). 3) glutamate metabolism; the pathway “alanine, aspartate and glutamate metabolism” was significantly regulated in NS (Holm test; p=0.028; ER=7.63), but not in LBN (Holm test; p=0.33; raw p-value=0.0042; ER=5.79). The “arginine biosynthesis” did not reveal significant differences in NS (Holm test; p=0.28; raw p-value=0.0040; ER=9.17) or LBN (Holm test; p=0.33; raw p-value=0.0042; ER=8.67). Panel A-C represent female NS (n=10, female LBN (n=10), male NS (n=10), male LBN (n=11).

Therefore, further analysis investigated the metabolic regulation on sex between males and females and were subsequently compared between conditions, NS and LBN exposure. A significant regulation of metabolites in the sex comparison (male vs female) was observed in the NS condition HILIC negative mode (49 significant features; corresponding to 33 significant validated metabolites) and HILIC positive mode (85 significant features; corresponding to 46 significant validated metabolites). In addition, the regulation of metabolites comparing sex (male vs female) in the LBN condition also showed a significant regulation in HILIC negative mode (51 significant features; corresponding to 36 significant validated metabolites) and HILIC positive mode (81 significant features; corresponding to 42 significant validated metabolites) (figure 4A).

The different HILIC settings (negative and positive mode) can reveal partially different metabolites and were merged to obtain a complete list of significantly regulated metabolites (NS: 63 and LBN: 61 merged and significant validated metabolites). The set of regulated metabolites in the male vs female comparison between NS and LBN conditions showed that 34.67% of the metabolites were specifically regulated in only NS or only LBN conditions (18.67%: 14 metabolites in NS and 16%: 12 metabolites in LBN of the total 75 metabolites) (figure 4B). These results point to the fact that there might be specific regulated pathways underlying these differentially regulated metabolites between stress conditions while comparing male vs female.

Next, a Metabolite Set Enrichment Analysis (MSEA) was performed, which aims to understand the biological significance of the enriched sets of metabolites, associated with specific biological pathways or processes between sexes comparing NS and LBN conditions. The MSEA revealed several pathways of interest when comparing between the NS and LBN conditions in relation to stress; 1) the purine metabolism pathway, 2) the pyrimidine metabolism pathways, and 3) the glutamate metabolism pathways, as observed in the “arginine biosynthesis” and “alanine, aspartate and glutamate metabolism” pathways (figure 4C).

The purine metabolism pathway showed a significant regulation in NS male vs female, as well as in LBN male vs female (figure 4C). In addition, the comparison NS and LBN showed similar enrichment ratios indicating a differential regulation of the purine metabolism pathway between males and females, regardless of stress exposure. The pyrimidine metabolism pathway showed no regulation in NS male vs female, but did show a significant regulation in LBN male vs female, while the enrichment ratios were similar between conditions (figure 4C). The glutamate metabolism pathway showed a significant regulation in NS male vs female, but not LBN male vs female, specifically in the “alanine, aspartate and glutamate metabolism”, but not in “arginine biosynthesis” (figure 4C). These results indicate the differential regulation of the glutamate metabolism pathway between NS and LBN when comparing males and females.

## Discussion

Exposure to ELS increases vulnerability to stress-related disorders, such as PTSD. The prevalence of PTSD is strongly influenced by sex. Animal models investigating ELS, in particular by using the limited bedding and nesting (LBN) model, have provided increasing evidence that LBN exposure affects rodents in a sex-specific manner ^43–47^. However, the sex-dependent impacts of HPA-axis signaling, metabolomic pathway analysis in stress-related brain regions, and fear memory formation on ELS remain to be fully elucidated. The development of open-source markerless pose estimation tools ^66^ and subsequently unsupervised behavioral analysis tools has enabled in-depth analysis that can explore previously unknown behavioral patterns ^67^. This advancement is crucial for enhancing the understanding of the behavioral outcomes associated with stress-induced fear memory formation. The present study contributes to a better understanding of those mechanisms by investigating HPA axis signaling in the body and brain, along with metabolomic pathway alterations in the BLA and behavioral responses to LBN-induced fear memory formation in a sex- and time-dependent manner.

### LBN disrupts different facets of the HPA axis in a time- and sex-dependent manner

The LBN model has been extensively utilized to investigate the effects of chronic ELS exposure on both physiological and behavioral outcomes. A common hallmark of LBN exposure is the reduction of body weight at P09 ^51^, which was confirmed in the current study, highlighting that the LBN model has stress-dependent effects in both sexes ^68,69^. In addition, the long-term effect of LBN on body weight between sexes is more variable and seems to be dependent on the age of testing and potential additional challenges throughout adulthood. The current study showed that at 3 months of age, females show a more persistent LBN-related body weight reduction phenotype compared to males. Other studies shown a similar effect at two months of age ^68^, but Arp et al. did not find this sex-dependent difference in 4 months old animals ^70^, indicating that both males and females eventually recover, but females show a longer recovery period. An opposite effect was observed in another study at eight months of age, where males showed a stronger body weight reduction compared to females ^69^, this might be explained by the different adult stress events (e.g., glucose- and insulin tolerance tests) and indicates different vulnerability toward such adult stressors after LBN exposure between sexes.

ELS exposure has been linked to dysregulation of the HPA axis, which can lead to an increased vulnerability state of stress-related psychopathology ^21,22^. A well-established phenomenon is the elevated levels of morning CORT baseline in females compared to males ^71–73^. The current study replicates this phenomenon in adult mice regardless of stress exposure, as also recently reported by Brix et al. ^69^. Interestingly, it was further observed that this sex-dependent difference is already apparent at the early age of P09, highlighting that the sex-specific differences are apparent already at the end of ELS exposure. An earlier study on LBN exposure in males showed a significant LBN-induced increase at P09 for baseline CORT in mice ^51^, but the current study found an increase in CORT only in LBN females. This might be explained due to the low baseline levels of CORT in males, which therefore, might show a higher variance, as the absolute CORT values between conditions are smaller. Nonetheless, the high increase of baseline CORT in LBN-exposed females is a good proxy for stress exposure, and at P09 is indicated to be higher in females compared to males. Moreover, we observed an opposite effect for adrenal weight at adult age, which was significantly increased for LBN-exposed males, but not females. This is in line with earlier research, that showed a similar effect in males at 1 month of age ^70^, but at later stages in adulthood, namely 4 months and 8 months, the adrenals in males were back to the same size as the non-stressed condition ^51,69^. This indicates that the adrenal size is influenced in a time-dependent manner, in which males are taking longer to recover their adrenal size to baseline after LBN exposure. Another facet of the HPA-axis reactivity was investigated via gene expression changes of FKBP5 in the brain. Previous research has identified a particularly high expression of the FKBP5 gene in the BLA and HIP under baseline, which is confirmed in the current study ^38^. Furthermore, FKBP5 gene expression has been shown to increase in a stressor-dependent manner in the BLA and HIP ^38,74,75^, but the immediate and long-term FKBP5 gene expression changes in response to LBN have remained elusive. We show that FKBP5 gene expression was not changed by LBN exposure directly after the stress at P09 in both sexes but was upregulated specifically in the CA1 region of the dorsal HIP of adult LBN-exposed males, but not females. Marrero et al. 2019 ^39^ showed that the overexpression of human FKBP5 in the forebrain induces specific downstream molecular changes in the dorsal HIP in adult ELS animals using the maternal separation paradigm. This coincides with the current finding that specifically the dorsal HIP shows upregulated FKBP5 expression and points to an altered molecular pathway mechanism after LBN exposure. In conclusion, we highlight a differential impact of LBN exposure across sexes. The immediate effects of LBN exposure at P09 are more pronounced in females, while interestingly, the prolonged effects of a dysregulated HPA-axis in adult age are affected exclusively in males.

### Fear acquisition is differentially affected by LBN across sexes

The formation of fear memory is a crucial aspect of understanding the underlying mechanism of PTSD. The specific alterations of anxiety and fear behavior can be investigated using animal models of fear conditioning. In line with previous research, we show that exposure to LBN reduces the fear response by lowering freezing behavior during both contextual- and auditory fear retrieval in males ^49^. In addition, we show that LBN exposure in females shows a similar reduction in freezing behavior during the contextual fear retrieval, but not in the auditory fear retrieval. Previous research has shown that LBN is linked to reduced synaptic plasticity markers within the dorsal HIP, which could explain the LBN-induced differences in fear retrieval by altering fear memory formation ^49^. However, the current study highlights an alternative explanation by specifically exploring the fear behavior during the acquisition of fear conditioning. Specifically, it was observed that the reduction in freezing behavior can already be found during the acquisition of the fear memory, in which LBN-exposed males show reduced freezing during the acquisition tones, and the females during the acquisition ITIs. A similar effect has been observed in other studies for male data, in which it was shown that the freezing directly after the fear acquisition is already lowered in LBN-exposed animals ^49,70,76^. Therefore, the difference in freezing during the retrieval phase is not only explained by differences in fear memory formation but also by an altered response at fear acquisition.

In addition, several studies have highlighted the relevance of distinguishing between different types of fear behaviors ^77–79^. The analysis of the behavioral data, using an unsupervised analysis, provides a promising way to explore novel behavioral patterns related to fear acquisition without prior behavioral categorization. This allows the exploration of the behavioral repertoire in a hypothesis-generating way, which can lead to the identification of novel behaviors within the specific methodological context ^80^. To further understand the sex-dependent fear behavioral differences during fear acquisition, the DeepOF open-source python package was deployed to perform an unsupervised analysis pipeline, which maps the representations of different fear behavior-related syllables across the different stages (tones and ITIs), conditions (NS vs LBN) and sex (female vs male) without any prior label information. Different fear-related behaviors were observed that were particularly altered in female, but not male mice. Interestingly, it was observed that specific behavioral clusters (e.g., cluster 0) are elevated in LBN-exposed females, which indicated a behavior related to the exploration of the environment. Conversely, other behavioral clusters (e.g., cluster 6) are reduced in LBN-exposed females, which were related to freezing behavior. Intriguingly, the observed behavior in “cluster 0” coincides with a previously identified active fear behavioral response, called “darting”, in which rapid locomotive movements are detected in primarily female rats ^79^. The behavioral syllables from “cluster 0” can also be allocated to an active fear behavioral response, but under non-stressed conditions are expressed in both females as well as males. However, the increased amount of the expression of “cluster 0” after LBN exposure is exclusively observed in females, which does indicate a sex-dependent effect on the active fear response.

### Sex and LBN exposure alter the purine, pyrimidine, and glutamate metabolism pathways in the BLA

The investigation of cellular metabolism in the BLA and the subsequent downstream analysis of metabolic pathways by MSEA are important tools to contribute to the understanding of the underlying neurobiological mechanisms related to sex differences of LBN exposure. A strong differential metabolic regulation in the BLA was found between sexes, but not stress condition. This was further highlighted by MSEA pathway analysis, which revealed altered purine-, pyrimidine-, and glutamate metabolism pathways between the sexes. These results highlight, for the first time, a striking role for metabolic regulation in the BLA on the sex-dependent differences between males and females. In addition, the differential metabolic profiles between sexes could also contribute to the explanation of the differential behavioral responses after LBN exposure. This was highlighted, for example, in the glutamate metabolism pathway, which showed a stronger altered regulation between males and females in the NS condition compared to the LBN condition. A possible explanation could be that LBN exposure alters glutamate metabolism in a similar way in males and females, masking the strong differences that are observed under baseline conditions.

Intriguingly, a recent study showed that there is a disbalance in GABAergic and glutamatergic gene expression levels in the ventral hippocampal neurons after LBN exposure in male mice ^81^. Taking into account the strong connection between the HIP and the BLA after stress ^82–84^ and their similar expression of stress-responsive genes, such as GR, MR and FKBP5, it is possible that a similar disbalance could be observed in the BLA. This would suggest that beyond the differences in gene transcription, the current study shows that the metabolic pathways related to glutamate metabolism in the BLA are altered as well in a sex-dependent manner for LBN exposure. In addition, Kos et al., 2023 ^81^ further show that changing the brain-wide balance between glutamatergic vs GABAergic signaling in LBN-exposed mice restores the ELS-induced affected behaviors, highlighting the importance of the glutamate and GABA in ELS affectedness, taken into account the current study, it is possible that this rescue effect is not only mediated via the ventral hippocampus but also the BLA.

## Conclusion

Taken together, the current study shows a sex-specific effect of LBN exposure on dysregulation of the HPA-axis, in which the adrenal weight, baseline CORT levels, and FKBP5 expression in several stress-related brain regions, including the BLA and subregions of the dorsal HIP, are altered in a time-dependent manner. In addition, we show that specific aspects of fear-related behavior, including the passive fear behavioral response via freezing behavior, but also an active fear response, as identified using an unsupervised analysis, are altered by LBN exposure in a sex-specific manner. The additional fear-related behavior that is expressed in “cluster 0” is contributing to a better understanding of the sex-dependent effects of fear memory acquisition and might influence the expression of the freezing behavior during contextual as well as auditory fear retrieval. The DeepOF unsupervised analysis provides an additional layer to explore the fear-related behaviors without prior assumptions and therefore, allows for hypothesis-generating behavioral analysis, which ultimately can lead to a better understanding of the stress-induced behavioral phenotype. Moreover, the current study identifies, for the first time, altered purine, pyrimidine, and glutamate metabolism pathways in the BLA on sex and LBN exposure. The current study proposes a potential link between these metabolic alterations and the observed fear-related behavioral outcomes, emphasizing the need for a comprehensive understanding of the metabolomic mechanisms underlying LBN exposure. Overall, these findings highlight the importance of considering sex-specific metabolic responses in understanding the neurobiological mechanisms underlying stress-related disorders and offer potential avenues for targeted interventions.

## Acknowledgments

The authors thank Lisa Rudolph, Daniela Harbich, and Bianca Schmid for their technical assistance, and the DeepLabCut development team for creating and maintaining the DeepLabCut software.

## Author contributions

JB and MVS conceived the study. JB performed the experiments, LvD, LMB, SN, HY, SM, VK and MS assisted with the experiments. Metabolomics was performed by KK and analyzed by TB and NG. All other data was analyzed by JB and with the assistance of LM and BMM. JB wrote the first draft of the manuscript and all authors contributed to the final manuscript version.

## Supplementary Information

**Supplemental Figure 1.**
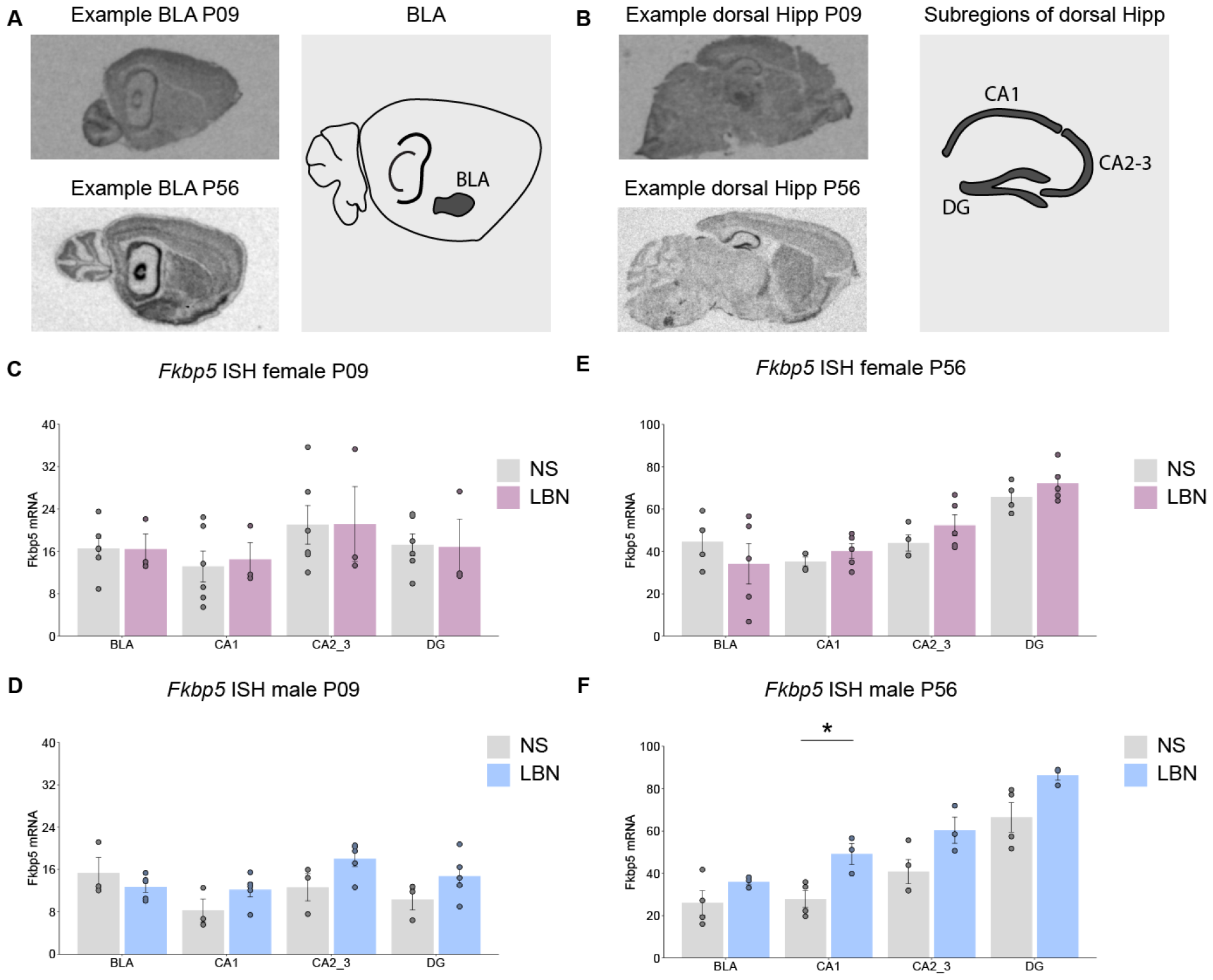
In-situ hybridization of *fkbp5* mRNA in the BLA and dorsal hippocampal subregions of non-stressed (NS) and stressed (LBN) mice. **A)** In-situ hybridization (ISH) scan of *fkbp5* mRNA expression at P09 and P56 in the BLA. **B)** In-situ hybridization scan of *fkbp5* mRNA expression at P09 and P56 in the subregions of the dorsal HIP. **C)** No differences were observed in females at P09 in the BLA (T(7)=0.03, p=0.98), CA1 (T(7)=-0.28, p=0.79), CA2-3 (Wx=11, p=0.71), and DG (T(7)=0.09, p=0.93). **D)** No differences were observed in males at P09 in the BLA (T(6)=1.03, p=0.34), CA1 (T(6)=-1.66, p=0.15), CA2-3 (T(6)=-1.98, p=0.096), and DG (T(6)=-1.49, p=0.19). **E)** No differences were observed in females at P56 in the BLA (T(7)=0.86, p=0.42), CA1 (T(7)=-1.14, p=0.29), CA2-3(T(7)=-1.27, p=0.25), and DG (T(7)=-1.21, p=0.27). **F)** A significant difference for elevated *fkbp5* mRNA expression was observed in the LBN condition for the males at P56 in the CA1 (T(5)=-3.38, p=0.020), but not in the BLA (T(5)=-1.44, p=0.21), CA2-3 (T(5)=-2.30, p=0.070), and DG (We(3.67)=-2.69, p=0.056). The bar graphs are presented as mean ± standard error of the mean and all individual samples as points. Panel C represent female P09 NS (n=6, female LBN (n=4). Panel D represent male P09 NS (n=3, male LBN (n=5). Panel E represent female P56 NS (n=4), female LBN (n=5). Panel F represent male P09 NS (n=4), male LBN (n=3).

**Supplemental Figure 2.**
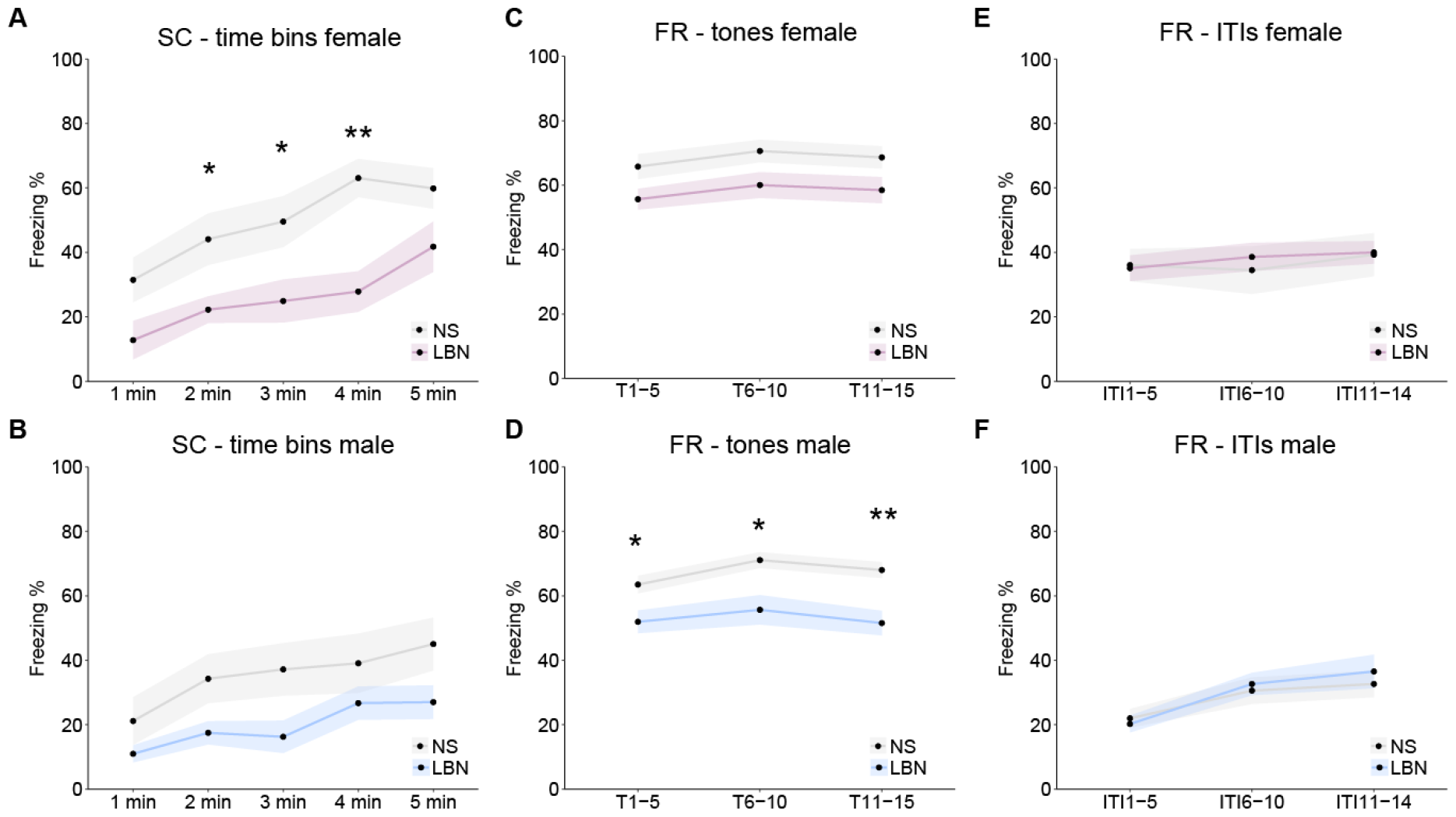
Time bin data for contextual and retrieval fear memory. **A)** The freezing behavior during 1min bins of the contextual fear memory in females. A significant main effect was observed using the two-way ANOVA on stress (F(1,85)=31.61, p<0.0001), and tones (F(4,85)=5.54, p=0.0005), but not on stress*tones (p=0.70). Post-hoc analysis using BH revealed a significant reduction of freezing in LBN females compared to NS for 2min (p=0.047), 3min (p=0.047), and 4min (p=0.004), but not 1min (p=0.069), and 5min (p=0.094). **B)** No significant main effect was observed using the Kruskal Wallis test (p>0.15). **C)** The freezing behavior during the auditory fear retrieval tones binned per 5 tones in females. A significant main effect was observed using the two-way ANOVA on stress (F(1,42)=11.59, p=0.001), but not for tones, or stress*tones (p>0.46). Post-hoc analysis using BH revealed no further significance between stress conditions (p=0.077). **D)** The freezing behavior during the auditory fear retrieval tones binned per 5 tones in males. A significant main effect was observed using the two-way ANOVA on stress (F(1,54)=27.96, p<0.0001), but not for tones, or stress*tones (p>0.24). Post-hoc analysis using BH revealed a significantly lowered freezing response in LBN males compared to NS at T1-5 (F(1,18)=6.695, p=0.019), T6-10 (F(1,18)=8.84, p=0.012, and T11-15 (F(1,18)=13.12, p=0.006). **E)** No significant main effect was observed using the two-way ANOVA for the fear retrieval ITIs in females (p>0.73). **F)** No significant differences were observed between LBN and NS males between the different ITIs in the fear retrieval task; the two-way ANOVA did reveal a significant main effect for ITIs (F(2.54)=6.80, p=0.002), but not for stress (F(1,54)=0.20, p=0.66), or stress*ITIs (F(2,54)=0.28, p=0.76). Further post-hoc analysis using BH revealed no significant differences (p>0.57). The timelines are presented as mean ± standard error of the mean and all individual samples as points. Panel A-F represent female NS (n=10, female LBN (n=10), male NS (10), male LBN (11).

**Supplemental Figure 3.**
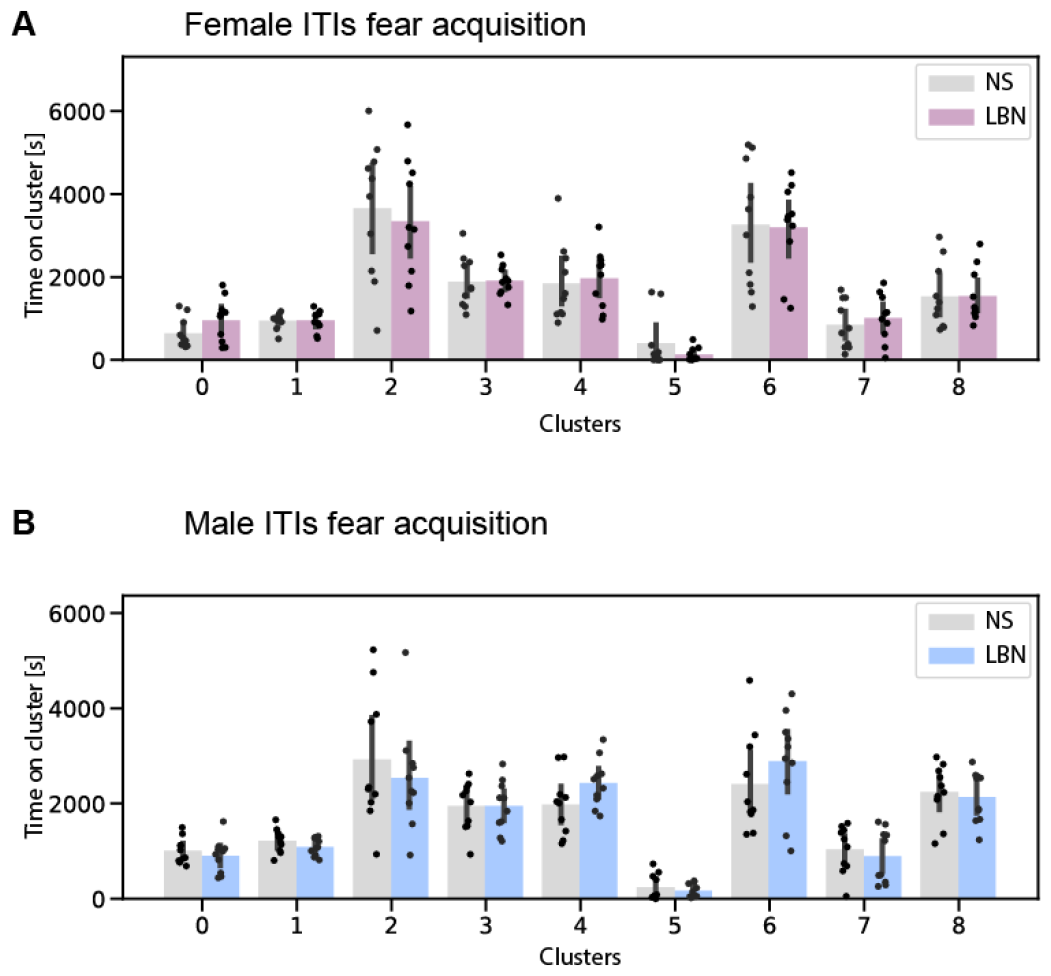
Unsupervised clusters during the ITIs. **A)** Cluster enrichment for the female fear acquisition data using all four ITIs. No significant differences were observed using the independent samples t-test corrected for multiple testing using Benjamini-Hochberg’s method across clusters (p>0.05). **B)** Cluster enrichment for the male fear acquisition data using all four ITIs. No significant differences were observed using the independent samples t-test corrected for multiple testing using Benjamini-Hochberg’s method across clusters (p>0.05). Bar graphs represent mean ± standard deviation of the time proportion spent on each cluster. The timelines are presented as mean ± standard error of the mean and all individual samples as points. Panel A-F represent female NS (n=10, female LBN (n=10), male NS (10), male LBN (11).

**Supplemental Figure 4.**
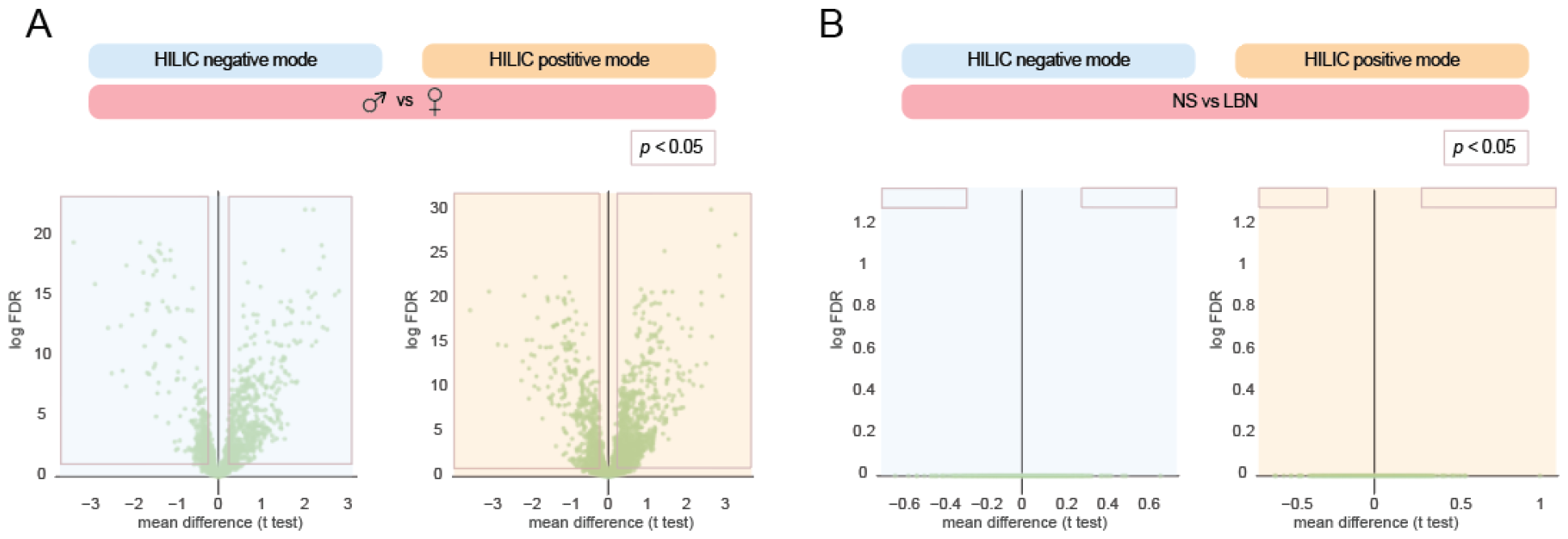
Metabolomics differential expression analysis on sex and stress condition. **A)** Volcano plots show a large differential expression profile comparing male vs females regardless of stress condition in HILIC negative and positive mode. **B)** Volcano plots show no differential expression profile comparing NS and LBN conditions, regardless of sex in HILIC negative and positive mode. Panel A-B represent female NS (n=10, female LBN (n=10), male NS (n=10), male LBN (n=11).

